# Causal gene regulatory network inference from Perturb-seq via adaptive instrumental variable modeling

**DOI:** 10.64898/2026.02.18.706642

**Authors:** Zhongxuan Sun, Hyunseung Kang, Sündüz Keleş

**Affiliations:** Department of Biostatistics and Medical Informatics, University of Wisconsin - Madison, 1205 University Ave, Madison, 53706, WI, USA; Department of Statistics, University of Wisconsin - Madison, 1205 University Ave, Madison, 53706, WI, USA

**Keywords:** Perturb-seq, instrumental variables, gene regulatory network

## Abstract

Inferring causal gene regulatory networks (GRNs) from observational single-cell data is challenging due to confounding. While Perturb-seq provides causal leverage, existing methods are often biased by heterogeneous CRISPRi knockdown efficiencies and restrictive assumptions like acyclicity. We present ADAPRE, a framework that treats CRISPR interventions as instrumental variables within a Poisson-lognormal model. By adaptively accounting for variable perturbation strength, ADAPRE recovers potentially cyclic structures and outperforms existing methods. Applied to a genome-wide K562 Perturb-seq dataset, it reconstructs networks enriched for known biological interactions and identifies coherent, leukemia-associated subnetworks, establishing a scalable approach for causal GRN inference.

## 1 Introduction

Gene regulatory networks (GRNs) represent how genes regulate one another’s expression. Traditional methods that rely on observational gene expression data struggle to deal with confounding and resolving complex feedback loops, both of which have been well-documented in human genetics [1]. The advent of pooled CRISPR screening with single-cell RNA-seq (Perturb-seq) [2–5] provides great opportunities to decipher GRNs by simultaneously perturbing thousands of genes with CRISPR technologies and quantifying expression changes at the single-cell level. One popular setting for Perturb-seq uses CRISPR intervention (CRISPRi), which employs an engineered guide RNA (gRNA) to direct a deactivated Cas9 enzyme to a specific gene, where it physically blocks transcription to reduce the gene’s expression rather than permanently cutting the DNA [6, 7]. The gRNA assignments are often regarded as random, providing opportunity to infer causal relations [3, 8].

Several computational frameworks leverage Perturb-seq for network reconstruction. DoTEARS [9] extends score-based learning to interventional data but assumes an acyclic structure, precluding the modeling of feedback loops. BICYCLE [10] can model steady-state dynamics with cycles, but it assumes perfect interventions. This makes it a poor fit for experiments using the partial and heterogeneous knockdowns produced by CRISPRi [6]. Another framework, LLCB [11], is also limited to CRISPR knockout data by its assumption of perfect interventions.

In contrast, *inspre* [12] uses a linear autoregressive structural equation model (SEM) which is a system of equations in which each gene’s expression is modeled as a function of other genes plus noise. In this framework, gRNA assignments are treated as instruments to estimate pairwise marginal average causal effects (ACEs) among genes. It then infers a sparse inverse of the ACE matrix to recover a cyclic direct-effect network, thereby addressing a key shortcoming of previous methods. However, *inspre* estimates pairwise ACE via two-stage least squares (2SLS) on control-relative, lane-wise z-scored expression (“internal z-normalization”), leaving the unique molecular identifier (UMI)-based molecule counting process unmodeled. With measurement model kept implicit, normalize-then-model workflows risk misattributing technical variation to expression and overlooking the Poisson mean–variance coupling of UMI counts [13]. Beyond this limitation, more importantly, we observe a perturbation-strength–dependent degree bias in the *inspre* framework: genes with stronger knockdowns are estimated to have higher out-degrees and are disproportionately inferred as hubs, distorting topology under heterogeneous CRISPRi strengths (Fig. 1a). Motivated by these limitations, we develop ADAPRE (ADAptive Penalized inverse REgression), which (i) explicitly models UMI counts with a Poisson–lognormal observation layer to disentangle measurement from expression and (ii) employs gene-specific adaptive penalties to correct strength-dependent degree bias in the estimated network. ADAPRE is a scalable and interpretable strategy for causal GRN inference from single-cell data with genome-scale CRISPRi perturbation, thereby bridging a critical gap between high-throughput perturbation assays and the reconstruction of the regulatory logic underlying cellular identity and disease.

**Fig 1.**
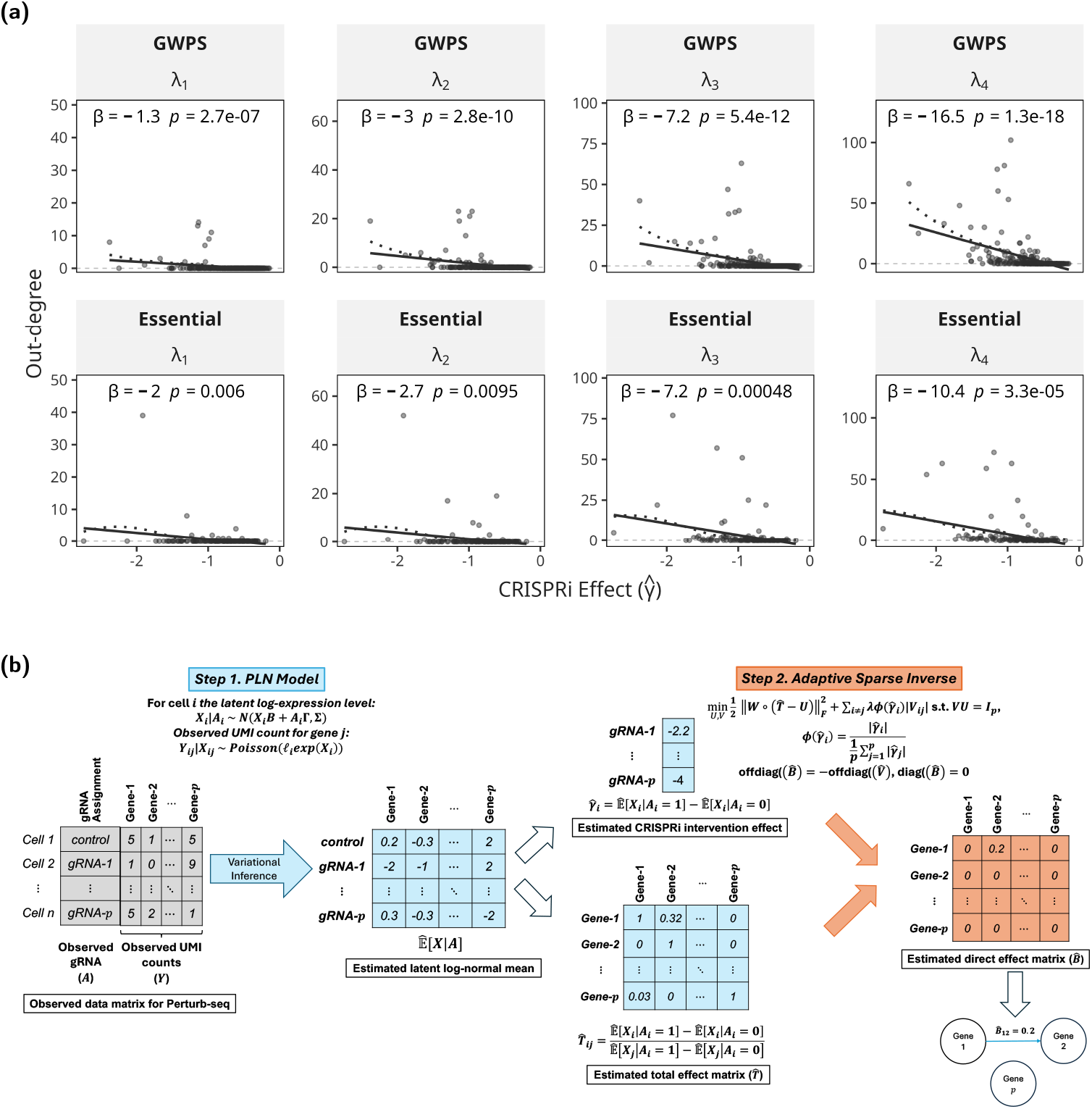
**(a) Perturbation–degree bias in *inspre* networks**.Scatterplots of estimated out-degree versus intervention strength 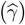, where 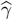 is the own-target CRISPRi knockdown effect measured in *z*-normalized expression units relative to control cells (more negative values indicate stronger knockdown). Rows correspond to datasets (GWPS vs. Essential); columns correspond to four decreasing L1-penalty strengths (*λ*_1_ > *λ*_2_ > *λ*_3_ > *λ*_4_), spanning very sparse ( ∼ 0.1%) to moderately sparse ( ∼ 3%) networks. Each point is a perturbed gene; dotted line is the LOESS curve; the solid line is the least-squares fit, with slope and *p*-value reported in panel headers. Across all settings, slopes are negative and significant (range − 16.55 to − 1.31; all *p* < 0.01), showing that genes with stronger perturbations are disproportionately inferred as high-degree regulators. This perturbation–degree bias indicates that *inspre* network topology can be distorted by heterogeneous intervention strength rather than true regulatory influence. **(b) Overview of the ADAPRE pipeline**. Starting from an *n* × *p* cell-by-gene Perturb-seq count matrix, ADAPRE fits a Poisson log–normal model to account for overdispersion and to estimate the average total effect of perturbing gene *i* on gene *j*. These estimates form a *p* × *p total effect matrix T* (diagonal fixed to 1). To separate direct from indirect pathways, ADAPRE applies adaptive, sparsity-penalized inverse regression to *T* to obtain a *p × p direct-effect matrix B*, interpreted as the signed, weighted GRN adjacency matrix.

## 2 Methods

### 2.1 Pervasive perturbation-degree bias in *inspre* networks

A key innovation of *inspre* is to turn interventional responses from large-scale CRISPRi Perturb-seq data into a direct-effect matrix 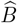 while allowing feedback loops (cycles) [12]. We begin by evaluating *inspre* estimated GRNs for potential biases. Applying *inspre* to estimate a 300-gene GRN 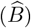 in K562 genome-wide CRISPRi Perturb-seq (GWPS) data from Replogle et al. [14] (after QC, *n* = 130,926 cells; *p* = 300 genes), we observed that genes with stronger CRISPRi knockdown (i.e., a stronger mean expression shift, denoted as *γ*) are disproportionately inferred as hubs across various levels of the tuning parameter that regularizes the network (Fig. 1a). Because CRISPRi efficacy is determined by guide design and locus context rather than gene function [6], a gene’s out-degree in the GRN should be independent of its perturbation strength. Therefore, the observed strength-degree dependence is likely an artifact of the estimation procedure.

To test whether the perturbation-degree bias generalizes beyond K562, we fitted GRNs on the CRISPRi Perturb-seq dataset of Schnitzler et al. [15], a large-scale perturbation study in human endothelial cells (teloHAECs) targeting 2,285 genes proximal to CAD GWAS loci. After QC, our analysis subset contains *n* = 29,097 cells across 212 gRNA groups (including a non-targeting control) targeting 211 genes; we retained 135 targets with significant knockdown (*p <* 0.05) and at least 100 perturbed cells per target for network fitting (Supplementary Table S2). Bias was quantified as the association between each gene’s out-degree and its own perturbation strength (knockdown magnitude). As shown in Fig. 4b (left column), similar to the trend observed in the K562 GRN, *inspre*-estimated GRN exhibits a strong perturbation-degree dependence, whereas incorporating adaptive shrinkage effectively removes this spurious coupling, yielding topology that is not driven by instrument strength (Fig. 4b - right column).

### 2.2 Overview of ADAPRE

ADAPRE is an end-to-end, two-stage, perturbation-aware framework for estimating causal gene regulatory networks (GRNs) from Perturb-seq. In stage 1, we estimate a matrix of total regulatory effects 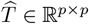, where 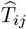 denotes the total (path-sum) regulatory effect of regulator *i* on target *j* under a linear SEM, from single cell UMI counts using a Poisson log-normal (PLN) model [16, 17]. Here, we set 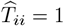. In stage 2, we recover the direct effect matrix *B* ∈ ℝ^*p×p*^ by solving a sparse inverse matrix problem that exploits the identity offdiag(*B*) = −offdiag(*T* ^−1^). Notably, in the inversion problem, there is an adaptive, row-specific *ℓ*_1_ penalty based on empirical instrument, i.e., knockdown, strength. The resulting output of ADAPRE is a sequence of sparse, directed-effect matrices 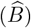 that represent estimated GRNs across a range of penalty levels. Subsequent sections describe ADAPRE in detail.

### 2.3 Model, assumptions, and identification

#### Observed data and notation

Suppose we observe a single-cell Perturb-seq experiment on *n* cells and *p* genes. Specifically, for cell *c* ∈ {1, …,*n*} and gene *i* ∈ {1, …,*p*}, we observe the raw UMI count, *Y*_*ci*_ ∈ ℕ_0_. This value represents the number of unique mRNA transcripts of gene *i* captured from cell *c*, where higher count indicates greater expression. The library size of the cell *c*, denoted by *ℓ*_*c*_, represents the total UMI count for the cell. We let *A*_*c*_ = (*A*_*c*1_, …,*A*_*cp*_) ∈ {0, 1}^*p*^ encode which among the *p* genes were perturbed in cell *c*. In the datasets considered in this manuscript, there is only one perturbation per cell, i.e., 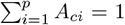 for perturbed cells *c*, and 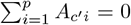 (i.e., *A*_*c*_′ = **0**) for control cells *c*^′^.

#### Modeling gene expression (PLN)

Count data from single-cell RNA-seq are often modeled with a negative binomial (NB) model to accommodate overdispersion [13]. However, despite advances in graphical-model discovery for count data [18, 19], extending the negative-binomial (Poisson–Gamma) model to a coherent multivariate graphical model is challenging: NB laws are not closed under the linear mixing implied by directed edges (and cycles), so nodewise NB conditionals generally do not assemble into a valid joint distribution. Given these limitations of NB models, we propose to use a Poisson log-normal model (PLN), first introduced by Aitchison and Ho [16], to model *Y*_*ci*_. PLN separates technical sampling (modeled with Poisson) from biological variability (modeled with the latent log-normal), consistent with the popular measurement-expression separation principle from Sarkar and Stephens [13]:

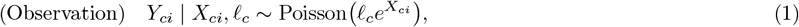

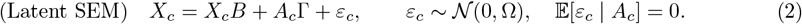

Here the observed count *Y*_*ci*_ depends only on the latent log-expression *X*_*ci*_ and the cell’s library size *ℓ*_*c*_, and is *conditionally independent* of the intervention *A*_*c*_ given (*X*_*ci*_, *ℓ*_*c*_). This encodes a measurement–expression separation: CRISPRi perturbs the biological state *X*_*c*_ via the SEM, while the Poisson sampling model for technical capture remains unchanged.

#### Structural SEM on latent expression

Similar to Brown et al. [12], we model the causal relation among the latent log-expressions using a linear, structural auto-regressive equation model (2). The matrix *B* ∈ ℝ^*p×p*^ encodes the gene regulatory network (GRN), with diag(*B*) = **0** ∈ ℝ^*p*^. Specifically, *B*_*ij*_ is the direct effect of gene *i* on gene *j*. We assume its spectral radius satisfies *ρ*(*B*) *<* 1, which guarantees that *I*_*p*_ − *B* is invertible (and hence *T* ≡ (*I* _*p*_ − *B*)^−1^ exists), and yields the convergent Neumann series 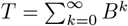. Thus 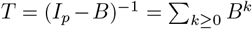 sums the weights of all paths, so *T*_*ij*_ is the total (direct + mediated) effect from *i* to *j*. The matrix Γ = diag(*γ*_1_,…, *γ*_*p*_) collects the direct CRISPRi effects on latent expression, with *γ*_*i*_ ≠ 0 for targeted genes and no off-diagonal entries; this structure encodes the relevance and exclusion assumptions for treating *A*_*ci*_ as an instrument for *X*_*ci*_.

#### Instrumental-variable assumptions and identification

To simplify notation, we now work at the population level for a randomly sampled cell and omit the explicit cell index. Let *X* = (*X*_1_, …, *X*_*p*_) denote the latent log-expression row vector and *A* = (*A*_1_, …, *A*_*p*_) the perturbation indicators. Under the low-MOI design, each cell is either unperturbed (*A* = **0**) or perturbed at exactly one gene, so *A*∈ { **0**, *e*_1_, …, *e*_*p*_}, where *e*_*i*_ is the *i*th canonical basis vector. Under the latent SEM model (2) and standard IV conditions (relevance, exclusion, independence, and no interference), we treat *A*_*i*_ as an instrumental variable for *X*_*i*_. The diagonal structure of Γ with nonzero entries *γ*_*i*_ encodes the relevance and exclusion assumptions, while we additionally assume that guide assignment is as-good-as random across cells and that each cell’s latent expression depends only on its own perturbation vector (independence and no interference).

Multiplying (2) on the right by (*I*_*p*_ − *B*) and taking expectations conditional on *A* yields

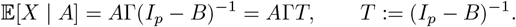

In particular, comparing cells perturbed at gene *i* (*A* = *e*_*i*_) to unperturbed controls (*A* = **0**) gives

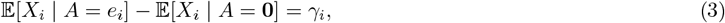

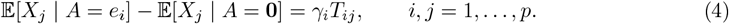

Equation (3) represents the own-gene knockdown effect of perturbing gene *i*, while equation (4) represents the mean shift in gene *j*’s latent expression when gene *i* is perturbed. Because Γ is diagonal, the effect of *A* on *X*_*j*_ must be mediated through the change in *X*_*i*_ and then propagated through the total-effect matrix *T*. Hence the Wald ratio [20] for outcome gene *j* with instrument gene *i* identifies the (*i, j*) entry of *T*:

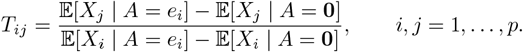

Given *T* = (*I*_*p*_ − *B*)^−1^, we have *T*^−1^ = *I*_*p*_ − *B*, so

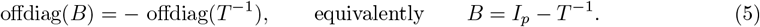

Thus, under the IV assumptions above and *ρ*(*B*) *<* 1, the off-diagonal entries of the direct-effect matrix *B* are identified from the joint distribution of (*X, A*) via the total-effect matrix *T*.

### 2.4 Estimation Procedure

#### Estimating latent means

Unlike *inspre*, which estimates the total-effect matrix *T* by two-stage least squares on *z*-normalized expression, we utilize the PLN framework to estimate latent expression shifts directly. We estimate the population-level latent mean expression for each perturbation group by fitting separate, intercept-only PLN models using the `PLNmodels` package [21]. Specifically, for each regulator gene *i* (defining the perturbation group *A* = *e*_*i*_) and for the control group (*A* = **0**), we fit an intercept-only PLN model to the corresponding subset of cells. The resulting group-specific intercept parameters provide estimates of the latent means, which we denote by 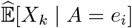 for cells perturbed at gene *i* and 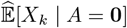 for unperturbed controls, for any gene *k*.

#### Role of the latent noise covariance Ω

In Eq. (2), Ω governs residual variation in the latent SEM and therefore affects the conditional covariance of latent expression, Cov(*X*|*A*) = *T*^⊤^Ω*T*, but it does not affect the conditional mean 𝔼 [*X* | *A*] = *A*Γ*T* since 𝔼 [*ε*_*c*_ | *A*_*c*_] = 0. Our estimator of total effects in Eq. (6) is mean-based, so we do not attempt to estimate Ω explicitly. Instead, the intercept-only PLN fits provide group-specific latent mean estimates while accommodating overdispersion and latent variability through nuisance dispersion parameters; we use only the fitted intercepts to construct 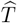. Standard errors 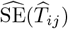 are obtained by bootstrap resampling of cells, which empirically propagates latent variability - including that induced by Ω - without requiring a plug-in 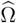.

#### Estimating total effects

We compute pairwise total effects using a Wald ratio based on differences in these estimated latent means. For each regulator *i* and target *j*, the estimated total effect is the shift in the target’s latent mean normalized by the regulator’s own knockdown strength:

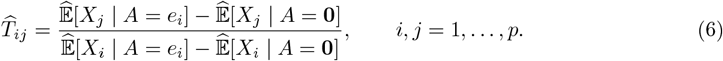

Here 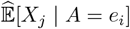 denotes the estimated latent mean of gene *j* from the PLN fit restricted to cells with perturbation vector *A* = *e*_*i*_ (and analogously for *A* = **0**, the unperturbed controls). Collecting entries yields the matrix 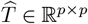, where we fix the diagonal 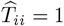. We obtain entrywise uncertainty 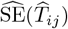 via a nonparametric bootstrap resampling of cells.

#### Estimating direct effects

By Eq. (5), the total-effect matrix satisfies *T* = (*I*_*p*_ −*B*)^−1^, so recovering the direct-effect network reduces to estimating *T*^−1^ via *B* = *I*_*p*_ − *T*^−1^. Because 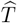 is noisy and GRNs are sparse, we estimate a denoised 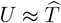 and a sparse left-inverse *V* ≈ *T*^−1^ (enforced by *V U* = *I*_*p*_) by solving

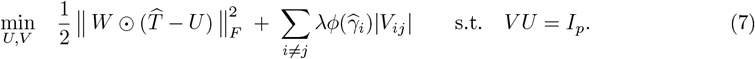

The matrix *W* is an entrywise weight matrix with 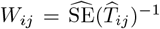, and 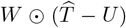 denotes the Hadamard product of two matrices. The terms 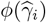 are *row-specific* weights of the form 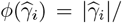 mean 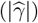, designed to adapt the sparsity of the matrix inverse to the row-specific perturbation strength. Intuitively, regulators with stronger instruments (larger 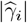) would otherwise dominate and be estimated as hubs; up-weighting their penalties counteracts this tendency, while weaker instruments are relatively down-weighted. When 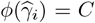 for all *i*, equation (7) reduces to the *inspre* estimator [12]. In summary, the optimization problem in equation (7) generalizes the *inspre* objective to allow sparsity patterns that adapt to instrument/perturbation strength.

#### Numerical optimization and convergence diagnostics

Numerically, we solve (7) using the alternating direction method of multipliers (ADMM) [22]. At each outer iteration, we minimize the augmented Lagrangian alternatingly in *U* and *V*: the *U*-update admits a closed form, while the *V* - update reduces to an *ℓ*_1_-penalized regression (lasso); dual variables are then updated. The outer-loop termination follows the *inspre* implementation based on relative objective stabilization, with early termination when the objective increases persistently, and a conservative iteration cap of `its`=300. Within each outer iteration, the lasso subproblems are solved by coordinate descent with inner budget `solve_its`=300, calibrated against a high-accuracy `solve_its`=400 reference. Convergence diagnostics, including constraint residuals 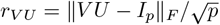, are reported in Supplementary Note S2 (Supplementary Table S1; Supplementary Figs. S2–S3).

## 3 Results

### 3.1 Simulation

#### Ablation of adaptive penalty

To isolate the contribution of *adaptive penalization*, independently of the count/PLN layer, we simulated Perturb-seq data from linear Gaussian SEMs (2), following settings similar to Brown et al. [12]. Starting from a direct-effect matrix *B* ∈ ℝ^*p×p*^ with diag(*B*) = 0, we enforced *ρ*(*B*) *<* 1 so that *T* = (*I*_*p*_ − *B*)^−1^ exists, allowing feedback cycles. Nonzero *B*_*ij*_ were drawn i.i.d. from a PERT distribution [23]. For each gene *j* ∈ {1, …, *p*}, we generated *N*_cont_ control cells and *N*_int,*j*_ intervention cells, with interventions entering as single-target exogenous perturbations of magnitude *γ*_*j*_. To induce heterogeneous instrument strength, we selected a subset *J*_strong_ with |*J*_strong_| = *p/*2 and set *γ*_*j*_ = 2*γ*_weak_ for *j* ∈ *J*_strong_ and *γ*_*j*_ = *γ*_weak_ otherwise, so that half of the regulators received a twofold stronger perturbation (colored separately in Fig. 2a).

**Fig 2.**
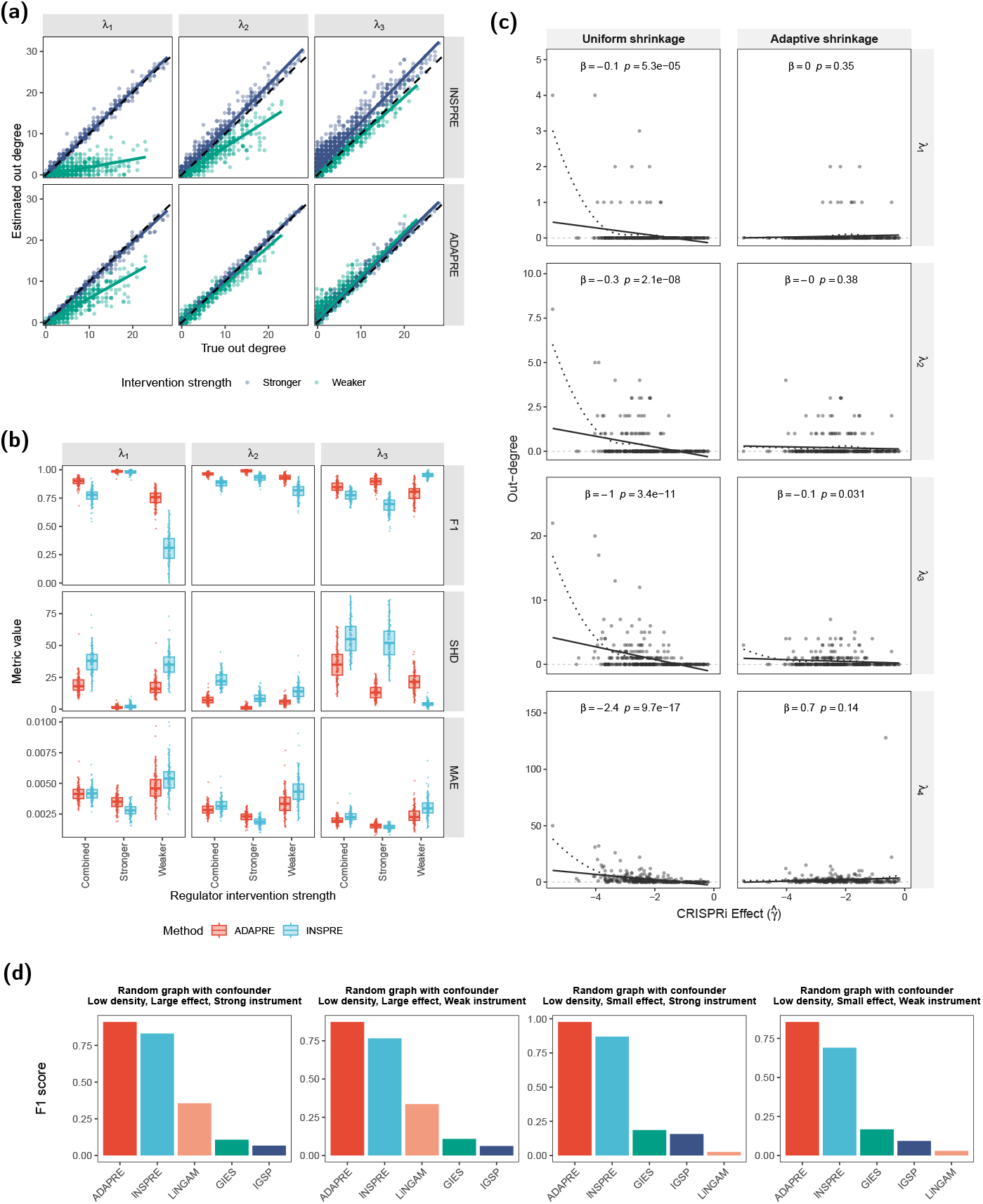
**(a) Degree calibration under heterogeneous interventions**. Each point is a gene; colors denote regulator strength; the dashed line is *y* = *x* and solid lines are group-wise linear regression fits. Row facet labels indicate the penalty scheme (*Uniform shrinkage (inspre)* vs. *Adaptive shrinkage (ADAPRE)*). **(b) Graph recovery across penalties and strength strata**. Boxplots of F1 (top), SHD (middle), and MAE (bottom) for combined, stronger-only, and weaker-only regulator sets, evaluated at *λ*_1_–*λ*_3_. **(c) Association between regulator out-degree and perturbation strength**. Association between regulator out-degree (*y*-axis) and CRISPRi perturbation effect size (*x*-axis): each point is a gene; the solid red line is a smooth fit with 95% confidence band. Under fixed shrinkage, stronger interventions spuriously inflate out-degree estimates, whereas adaptive shrinkage eliminates this dependence across penalty levels. **(d) Benchmarking against state-of-the-art methods**. F1 scores on random graphs with confounders. ADAPRE (red) outperforms baselines including *inspre* (blue) and other methods (LiNGAM, GIES, IGSP), and maintains high accuracy even in weak instrument settings (right panels) where alternative methods fail.

Recovery of *B* was summarized by structural Hamming distance (SHD), precision, recall, and F1 across a 20-point penalty grid, with sparsity matched between methods. For visualization, we report three representative penalties (*λ*_1_, *λ*_2_, *λ*_3_) chosen near the peak F1 for each method (Fig. 2b), and we stratify all metrics by regulator strength (genes in *J*_strong_ versus its complement). As shown in Fig. 2a, adaptive penalization largely removes the instrument-strength-degree bias: for both “stronger” and “weaker” regulators, estimated out-degree tracks the identity line and the group-specific regression slopes converge toward one another. This calibration translates into improved network recovery (Fig. 2b): across representative penalties, ADAPRE attains higher F1 and lower SHD and mean absolute error (MAE), with the largest gains for weaker regulators.

#### Benchmarking against additional baselines

Having established the effect of adaptive penalization in isolation, we compared ADAPRE with the baselines assessed in *inspre*, including the observational-only LiNGAM [24] and the interventional algorithms GIES [25] and IGSP [26]. We evaluated performance on random graphs with cycles and a latent confounder, across regimes varying in graph density (low/high), regulatory effect size (small/large), and instrument strength (strong/weak).

Fig. 2d displays F1 scores for these under cyclic, confounded regimes. Across all combinations of density, effect size, and instrument strength, ADAPRE consistently achieves the best performance, maintaining high F1 even under weak instruments and small effects. *inspre* typically ranks second, whereas LiNGAM, GIES, and IGSP lag substantially behind in these cyclic, confounded settings. This ordering, i.e., interventional methods outperforming observational ones, with ADAPRE further improving on *inspre*, is consistent with the benchmarking hierarchy reported by Brown et al. [12], while highlighting the additional gains provided by strength-adaptive penalization when perturbation efficacy is heterogeneous.

#### Computational scalability

To evaluate scalability, we profiled runtime and memory usage as a function of number of genes *p*, decomposing costs into Stage 1 (preprocessing and total-effect estimation, including PLN preprocessing when used) and Stage 2 (sparse inversion). We found a distinct trade-off: sparse inversion is the primary wall-clock bottleneck, whereas Stage 1 dictates the peak memory footprint (RSS) due to cell-by-gene data handling (Supplementary Note S1; Fig. S1).

### 3.2 Empirical validation on real-world Perturb-seq data

#### ADAPRE corrects perturbation–degree bias

On the K562 genome-wide Perturb-seq dataset [14], we plotted inferred out-degree versus CRISPRi effect size 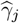 across four sparsity-matched penalties (*λ*_1_ – *λ*_4_) (Fig. 2c). With the PLN layer and total-effect construction held fixed, uniform penalties show a clear strength–degree bias (more negative 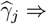 larger out-degree), whereas ADAPRE yields slopes near zero, indicating degree estimates independent of intervention strength. Correction is also evident in the teloHAECs data, as shown in Fig. 4b.

#### Validation against orthogonal datasets

We benchmarked enrichment of inferred edges against curated resources across 20 sparsity levels (Fig. 3a). **CORUM** [27] and **STRING** [28] provide *undirected* gene-gene relationships: we connected gene pairs co-annotated to a protein complex (CORUM) or recorded as protein-protein interactions (STRING), with a *STRING (physical)* subset for direct interactions. Because these are undirected, we symmetrize the estimate, counting a pair {*i, j*} as present if either *i*→ *j* or *j* → *i* is nonzero. For TF binding from ChIP–seq data [29], we define a *directed* TF→target reference by intersecting TF peak calls with (i) promoter windows (e.g., ±1kb around annotated TSS; “promoter”), (ii) enhancer annotations from EnhancerAtlas v2.0 (“enhancer”) [30], and (iii) their union. Orientation is assigned from the TF to the gene, and candidate regulators are restricted to TF genes. To complement enrichment-based validation, we additionally summarize directed TF→target accuracy using the baseline-adjusted macro-precision ΔPrec@50 (Variant B), which quantifies top-50 precision beyond the TF-specific reference density (Supplementary Note S3; Supplementary Fig. S4).

**Fig 3.**
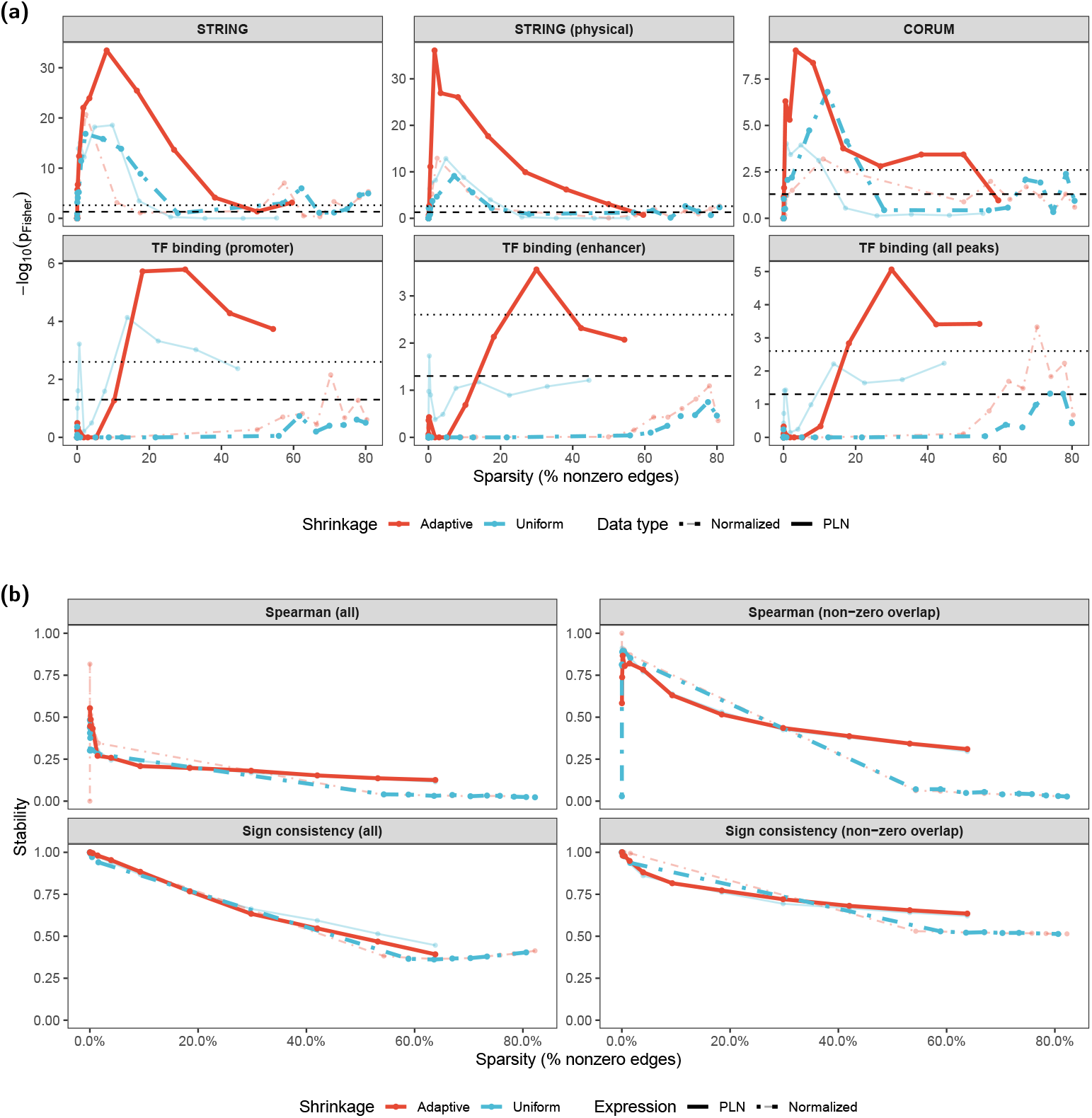
**(a) External validation against independent references**. Enrichment of inferred edges as a function of network sparsity, quantified by Fisher’s exact test ( −log_10_ *p*) against STRING (all associations), STRING (physical-only), CORUM complexes, and TF–target binding (ChIP–seq; promoter, enhancer-linked, union). Horizontal lines mark nominal 0.05 and Bonferroni-corrected 0.05*/*20 thresholds. Line color encodes shrinkage scheme (red = adaptive, blue = uniform); linetype encodes expression layer (solid = PLN, dashed = normalized expression). **(b) Split-half stability of estimated networks**. Stability of edge weights across random 50–50 splits as a function of sparsity, assessed using Spearman correlation and sign consistency on all edges and on the nonzero overlap. Stability decreases with density, but overlap-restricted metrics remain high and adaptive and uniform variants perform comparably.

At each sparsity level and for each resource, we formed a 2 *×* 2 contingency table over the appropriate edge universe (undirected pairs or directed TF→target edges) with entries: 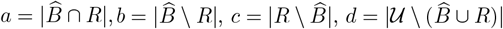. Enrichment is assessed by a one-sided Fisher’s exact test with alternative OR > 1, and we report − log_10_ *p* (Fig. 3a). Across CORUM, STRING (all/physical), and TF binding references, ADAPRE with a PLN expression layer yields stronger and more sustained enrichment along the sparsity path, consistent with benefits from both count-aware modeling and adaptive penalization.

#### Stability of estimated GRN

To test whether PLN modeling and adaptive penalization affect network stability, we split cells into two independent halves stratified by intervention target and reconstructed GRNs under all four method combinations (PLN vs. *z*-score; adaptive vs. uniform). For each method, we matched the two split-specific networks by sparsity level and quantified agreement using (i) the Spearman correlation of edge weights between the estimated 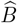 matrices and (ii) the sign-consistency rate on the *overlapping* set of nonzero edges. As shown in Fig. 3b, stability is largely unchanged across modeling choices and is instead driven by the sparsity level along the path.

#### Replication across datasets

To evaluate the replicability of estimated GRNs from GWPS, we leveraged the second Perturb-seq screen data from Replogle et al. [14], which harbored 2,057 common essential genes sampled 6 days after transduction (the “Essential” dataset; after QC, *n* = 45,222 cells). We filtered the 300-gene list to a subset of 134 genes perturbed in both datasets. We then separately estimated GRNs for the 134 genes using (i) GWPS data and (ii) Essential data. We matched the networks by sparsity and quantified agreement using (i) Spearman correlation of edge weights and (ii) the proportion of overlapping nonzero edges with consistent sign (Fig. 4a). Across the sparsity path, cross-dataset correlations were consistently positive (*ρ* ≈ 0.25–0.4 for all edges; up to ∼ 0.8 when restricted to overlapping nonzero edges), and sign agreement remained high (>80% at moderate sparsity). These results indicate that key directional effects inferred from the GWPS experiment are replicable in the Essential dataset, despite differences in sampling time.

**Fig 4.**
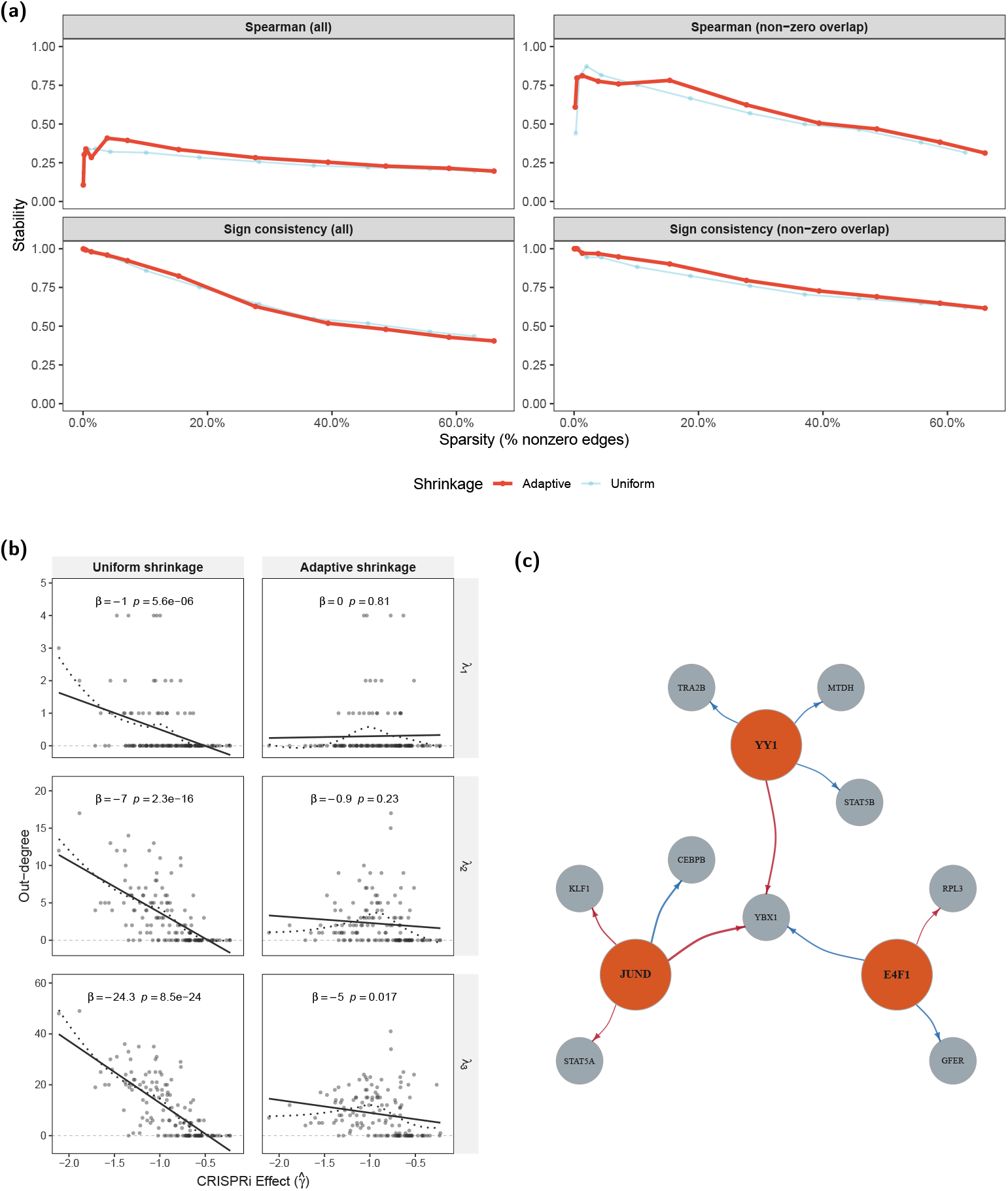
**(a) Cross-dataset reproducibility of inferred networks**. Cross-screen reproducibility between two independent K562 Perturb-seq datasets (GWPS vs. Essential): Spearman correlation of edge weights and sign consistency versus sparsity show persistent positive agreement overall and high concordance on the nonzero-overlap. **(b) Attenuation of instrument-strength bias in endothelial cells**. Endothelial (teloHAEC) Perturb-seq confirms the instrument-strength bias: under uniform shrinkage, estimated out-degree decreases with stronger CRISPRi effect (more negative 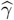), whereas adaptive shrinkage markedly attenuates this dependence across penalties. **(c) Functionally supported K562 regulatory modules**. Subnetworks centered on *YY1, JUND*, and *E4F1*, constructed by filtering the 10% edge-density K562 GRN for edges supported by curated TF binding at promoters/enhancers [30, 31] and STRING protein-association evidence [28]. Displayed targets contribute to the selected enriched term(s) for each hub. For readability, we show the top-*k* strongest outgoing edges per hub ranked by 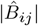 (with *k* = 4 except hubs with fewer eligible targets). Edge colors denote the sign of the estimated direct effect (blue: positive; red: negative). Modules highlight a *YY1*-centered RNA/gene-expression neighborhood, a *JUND*-centered stress-responsive and hematopoietic-linked neighborhood, and an *E4F1*-centered cytokine-response neighborhood, with *YBX1* appearing as a shared target in the displayed slice.

### 3.3 Perturbation-supported subnetworks in K562

We refined the ADAPRE K562 GRN (10% edge-density) by retaining directed regulator→target edges supported by curated TF binding [30, 31] and STRING protein associations [28]. Because STRING provides undirected association support, edge direction and sign were inherited from the perturbation-derived estimates. We assessed biological coherence via GO biological-process enrichment and pathway/hematopoietic signatures; given the restricted universe of expressed genes, we interpret these enrichments as contextual consistency rather than independent validation. For visualization (Fig. 4c), we show the top-*k* outgoing edges per hub among targets contributing to the selected enriched term(s) (with *k* = 4 except hubs with fewer eligible targets).

The resulting evidence-supported neighborhoods (Fig. 4c) reveal distinct functional patterns centered on *YY1, JUND*, and *E4F1*. The *YY1* neighborhood connects core expression machinery to signaling-adjacent effectors, showing positive direct-effect estimates on *TRA2B, MTDH*, and *STAT5B*, alongside a negative direct-effect estimate on *YBX1*. This structure is enriched for RNA biosynthetic and metabolic processes, consistent with YY1’s established role as a multifunctional chromatin regulator [32]. In contrast, while GO BP terms for the *JUND* neighborhood are generic, complementary enrichment in signature collections highlights stress-responsive (TNF/NF-*κ*B-related) and hematopoietic contexts. *JUND* shows a positive direct-effect estimate on *CEBPB* and negative direct-effect estimates on *KLF1* and *STAT5A* (Fig. 4c), placing it at the interface of myeloid/inflammatory regulation and erythroid nodes. Finally, the *E4F1* neighborhood exhibits mechanistic coherence with mitochondrial and translational homeostasis [33], driven by positive direct-effect estimates on *GFER* and *YBX1* and a negative direct-effect estimate on *RPL3*. Although the strict GO BP signal emphasizes cytokine responses, the target identities support prior evidence linking E4F1 to oxidative stress responses and leukemic cell survival [34]. Notably, *YBX1* emerges as a convergent integration point receiving opposing inputs (negative from *YY1* and *JUND*; positive from *E4F1*), nominating this antagonistic control axis for targeted experimental follow-up.

A direct comparison between ADAPRE and *inspre* GRNs at matched sparsity is provided in Supplementary Note S4 (Supplementary Fig. S5). This comparison shows that, even at matched sparsity, ADAPRE and *inspre* exhibit high signed Jaccard distance and only partial sign concordance among shared edges, indicating that adaptive penalization materially reshapes the inferred GRN rather than simply rescaling edge weights.

## 4 Discussion

ADAPRE addresses two major limitations in causal GRN estimation from CRISPRi Perturb-seq: (i) the absence of a probabilistic model for UMI counts that explicitly accounts for the technical sampling process and biological variation, and (ii) pervasive efficacy-driven biases that inflate the estimated out-degree of strongly knocked-down genes (Fig. 2a). By pairing a Poisson–lognormal model with instrument-strength–informed regularization, ADAPRE yields directed networks that effectively remove efficacy-driven bias (Fig. 2c), achieve better performance in both simulations and real-data applications (Fig. 2b; 3a), and preserve the stability and replicability of the estimates (Fig. 3b; 4a).

Several limitations suggest avenues for extension. First, identification relies on standard IV assumptions. CRISPRi may exhibit off-target or shared-pathway effects that violate exclusion [14], motivating more robust methods and sensitivity analyses. Second, we focus on single-perturbation designs; extending the estimator to multiplex perturbations (i.e., multiple knockdowns in one cell) will broaden applicability. Third, adaptive penalization depends on empirical instrument strength; more principled shrinkage (e.g., empirical-Bayes priors on rows of *B*) and joint tuning with biological priors (TF–target constraints) could further improve and stabilize estimation. Finally, advances in biotechnology are yielding large-scale multiome perturbation screens [35]; extending the method to leverage multimodal profiling will be important.

In sum, ADAPRE links a count-aware measurement model with adaptive IV-based structure learning to deliver scalable, interpretable GRNs from Perturb-seq. By identifying coherent regulatory hubs and narrowing the gap between high-throughput perturbation assays and mechanistic modeling, this framework advances our ability to decipher the complex wiring of the cell in both health and disease.

## 5 Code Availability

The source code for ADAPRE, together with a minimal test dataset and scripts to reproduce the simulation experiments in this paper, is publicly available at https://github.com/zsun263/DEMO_RECOMB_2026.

## 6 Data Availability

This study did not generate new sequencing data. We analyzed publicly available Perturb-seq datasets. Processed K562 genome-wide CRISPRi Perturb-seq data from Replogle et al. were obtained from the Figshare+ dataset (ID: `20029387`) [14]. The teloHAEC CRISPRi Perturb-seq dataset from Schnitzler et al. (PMID: 38326615) is available in GEO under accession `GSE210523` [15].

