## supplementary note for "Causal gene regulatory network inference from Perturb-seq via adaptive instrumental variable modeling"

### Supplementary Material

#### Supplementary Note S1: Computational scalability

We benchmarked runtime and memory as a function of number of genes $p$ and decomposed computation into (i) PLN preprocessing (ADAPRE stage 1), (ii) *inspre* end-to-end total-effect estimation via 2SLS followed by sparse inversion, and (iii) solver-only sparse inversion using fixed $(T, W)$ inputs exported from end-to-end runs (Fig. S1). Benchmarks were run on a Linux CPU node with two AMD EPYC 7513 processors (64 physical cores total) and 512 GB RAM; PLN used 64 process workers and inversion used 40 CPU cores. Across $p = 50$–$300$, sparse inversion dominates wall-clock time while stage 1 operations dominate memory, consistent with inversion operating on $p \times p$ summary matrices whereas preprocessing/total-effect estimation must handle cell-by-gene quantities. PLN results are additionally shown at $p = 500$ to illustrate stage 1 scaling (see caption for caveats).

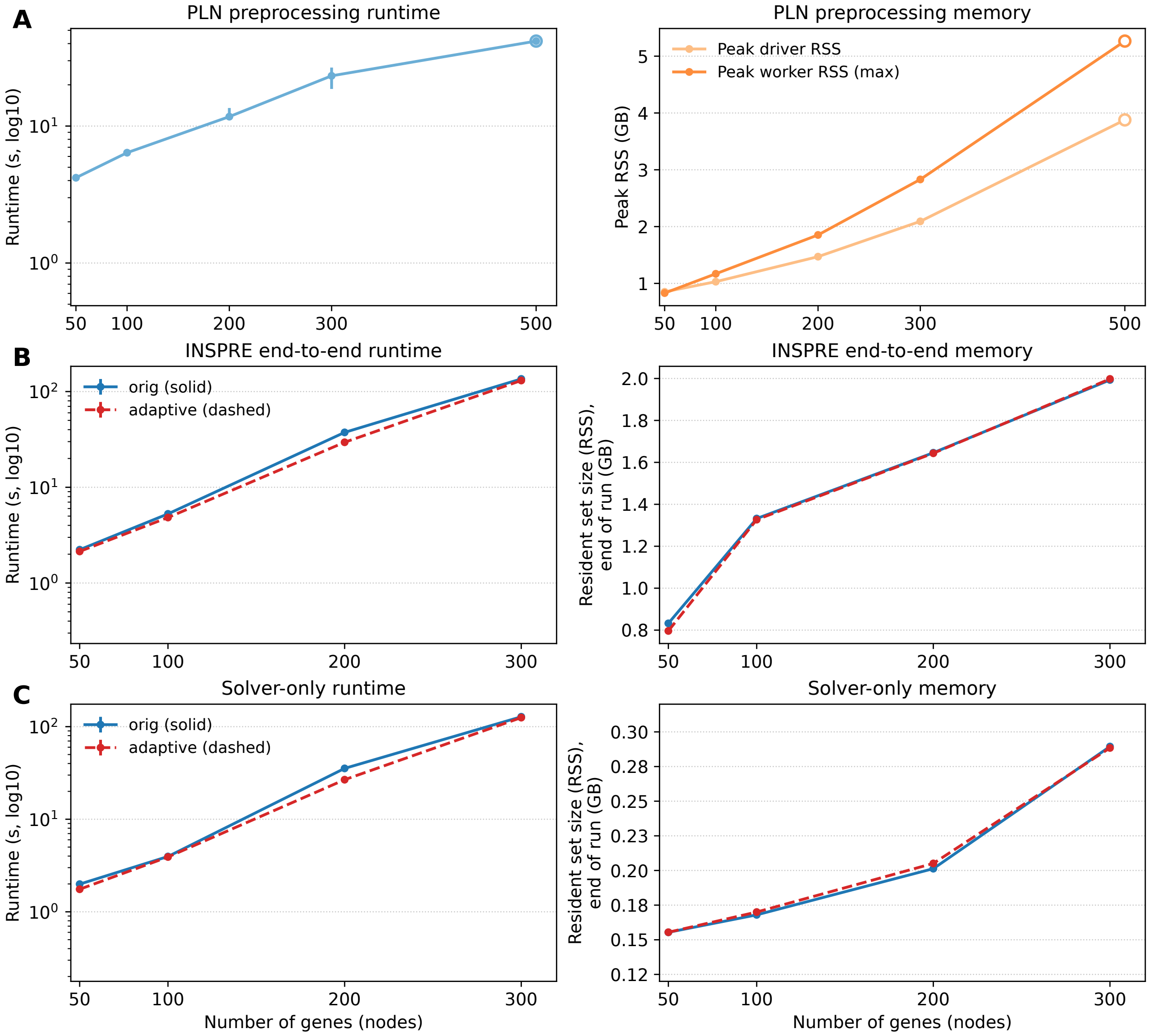


Figure S1: **Computational scaling of ADAPRE/*inspre* components with the number of genes $p$.** (A) PLN preprocessing runtime (log scale) and peak memory (peak RSS for the driver and maximum worker). (B) *inspre* end-to-end runtime and end-of-run memory ($RSS_{end}$) for 2SLS total effects + sparse inversion. (C) Solver-only sparse inversion runtime and $RSS_{end}$ using fixed $(T, W)$. Points denote medians with IQR error bars (PLN: seeds 1–50; *inspre*/solver: 5 repeats). PLN uses 64 workers; inversion uses 40 CPU cores. In (B–C), *orig* denotes the original (uniform-penalty) solver used by *inspre*, and *adaptive* denotes the same inversion backbone with our adaptive penalization. Panel (A) includes $p = 500$ to illustrate PLN scaling; this setting includes failed group fits and is therefore shown as a stress-test point rather than a clean all-success timing baseline.

#### Supplementary Note S2: Optimization convergence diagnostics

This note reports convergence diagnostics for the ADMM solver used for sparse inversion in Section 2.4, including the outer-loop termination rule, direct verification of the enforced constraint $VU = I_p$, empirical iteration counts across $\lambda$, and sensitivity to the inner lasso subproblem budget.

**Outer-loop termination rule.** Convergence of the outer ADMM loop is monitored using the augmented Lagrangian value $L^{(k)}$ at iteration $k$. Following the `inspre_worker()` implementation, convergence is declared when the relative change $|L^{(k)} - L^{(k-1)}|/L^{(k-1)}$ falls below `delta_target` $= 10^{-4}$ for two consecutive iterations. Early termination is triggered if $L^{(k)}$ increases for three consecutive iterations. All experiments additionally enforce a conservative maximum of `its` $= 300$ outer iterations. In all runs reported here, termination occurred by the stabilization criterion rather than the early-termination safeguard or the iteration cap.

**Constraint satisfaction.** We quantify satisfaction of the enforced constraint $VU = I_p$ at termination using the relative Frobenius residual $r_{VU} = \|VU - I_p\|_F/\sqrt{p}$. Table S1 summarizes these residuals over the full $\lambda$ grid for the GWPS dataset (`solve_its`=300). For completeness, we also report the auxiliary diagnostic $r_{UV} = \|UV - I_p\|_F/\sqrt{p}$; $UV = I_p$ is not enforced by the optimization formulation and can therefore deviate without violating feasibility of the fitted solution. Across settings, $r_{VU}$ remains small over the $\lambda$ grid (median $3 \times 10^{-3}$–$9 \times 10^{-3}$; maximum $\leq 7 \times 10^{-2}$), indicating that the enforced constraint $VU = I_p$ is well satisfied at ADMM termination.

Table S1: Constraint satisfaction at ADMM termination. Relative residuals for the enforced constraint $r_{VU} = \|VU - I_p\|_F/\sqrt{p}$ and the auxiliary diagnostic $r_{UV} = \|UV - I_p\|_F/\sqrt{p}$, summarized over the full $\lambda$ grid for the GWPS dataset with `solve_its`=300. We report $r_{UV}$ only as an auxiliary diagnostic; $UV = I_p$ is not enforced and can deviate without violating the optimization constraint.

| Model | Shrinkage | Method | `solve_its` | Median $r_{VU}$ | Max $r_{VU}$ | Median $r_{UV}$ | Max $r_{UV}$ |
|---|---|---|---|---|---|---|---|
| PLN | Uniform | *inspre* | 300 | 0.00530 | 0.0282 | 0.00552 | 0.0737 |
| PLN | Adaptive | ADAPRE | 300 | 0.00949 | 0.0531 | 0.00970 | 0.2109 |
| Normalized | Uniform | *inspre* | 300 | 0.00319 | 0.0699 | 0.00933 | 0.3064 |
| Normalized | Adaptive | ADAPRE | 300 | 0.00305 | 0.0298 | 0.00323 | 0.1288 |

**Outer ADMM convergence.** Figure S2 shows the number of outer ADMM iterations required to reach termination across $\lambda$ indices, stratified by dataset, model, and method. All runs terminate well below the conservative cap of `its` $= 300$ (red dashed line), indicating that reported solutions are determined by the stopping rule rather than truncation at the iteration limit.

**Sensitivity to the inner lasso budget.** We evaluate sensitivity to the inner coordinate-descent budget by comparing inferred sparsity patterns using `solve_its` $\in \{10, 50, 200, 300\}$ against a high-accuracy reference fit (`solve_its` $= 400$). Figure S3 shows that discrepancies decay rapidly as `solve_its` increases, and that `solve_its` $= 300$ is effectively indistinguishable from the reference across settings, supporting numerical stability of the inferred edge set at the chosen inner-solver budget.

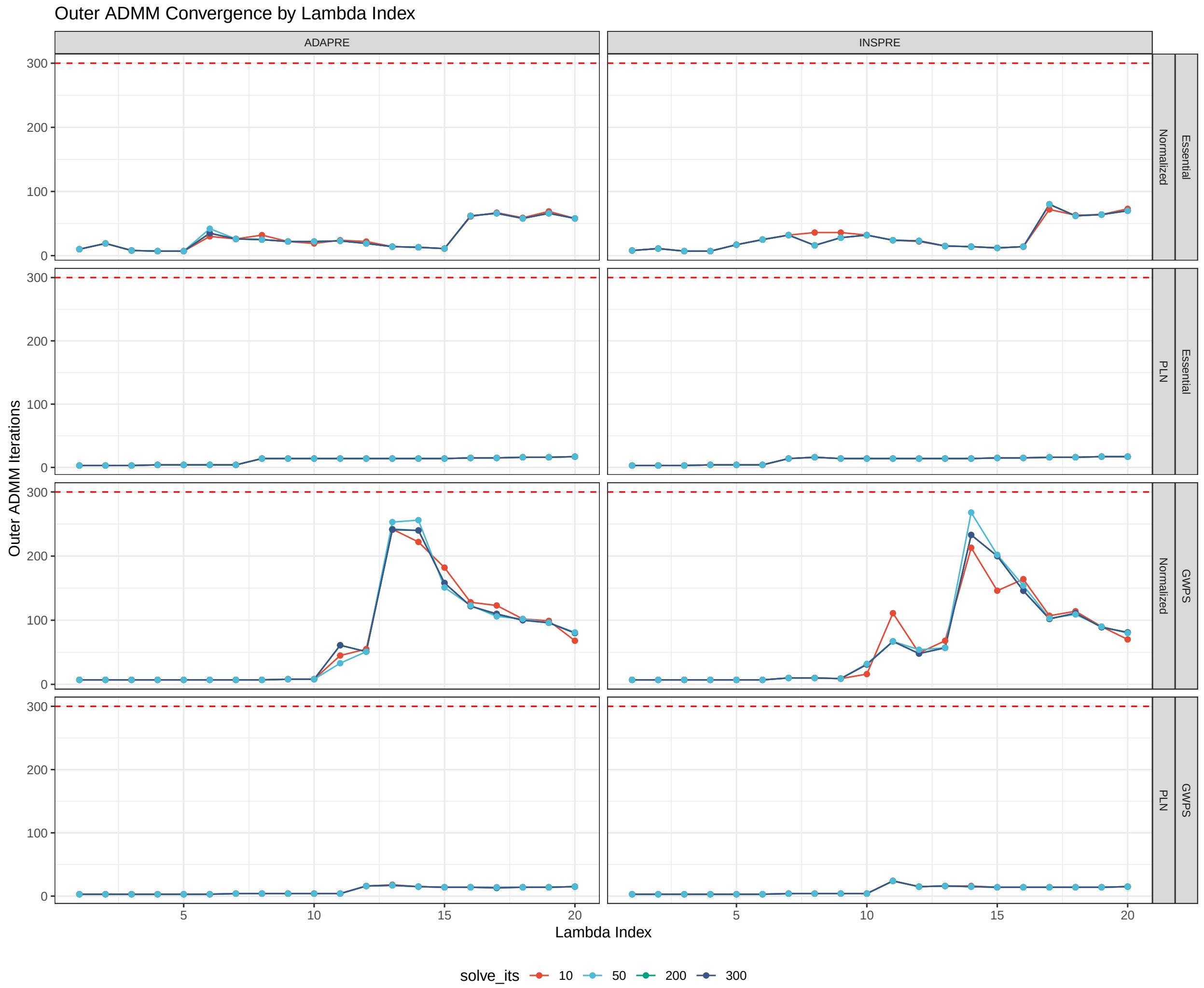


Figure S2: Outer ADMM convergence by $\lambda$ and inner-solver budget. Number of *outer* ADMM iterations until termination across $\lambda$ indices for ADAPRE and *inspre*, stratified by dataset (GWPS/Essential) and model (Normalized/PLN). Colors indicate the inner lasso coordinate-descent budget `solve_its` $\in \{10, 50, 200, 300\}$. The red dashed line marks the conservative cap `its` $= 300$; no run reaches this cap.

Table S2: Datasets used for model fitting after quality control and analysis subsetting. Cell counts ($n$) are reported after QC; $p$ denotes the number of genes used for network inference. Notes: all datasets include a non-targeting control group. GWPS uses a 300-gene panel for bias analyses and the K562 case study. Essential is the overlap subset used for cross-screen replication. CAD (teloHAEC) targets are retained by knockdown significance ($p < 0.05$) and $\geq 100$ cells per target (min 101; median 108.5).

| Dataset | Cell type | $n$ cells | $p$ genes | gRNA groups | targets used |
|---|---|---|---|---|---|
| GWPS | K562 | 130,926 | 300 | 301 | 300 |
| Essential | K562 | 45,222 | 134 | 135 | 134 |
| CAD (teloHAEC) | teloHAEC | 29,097 | 135 | 212 | 135 |

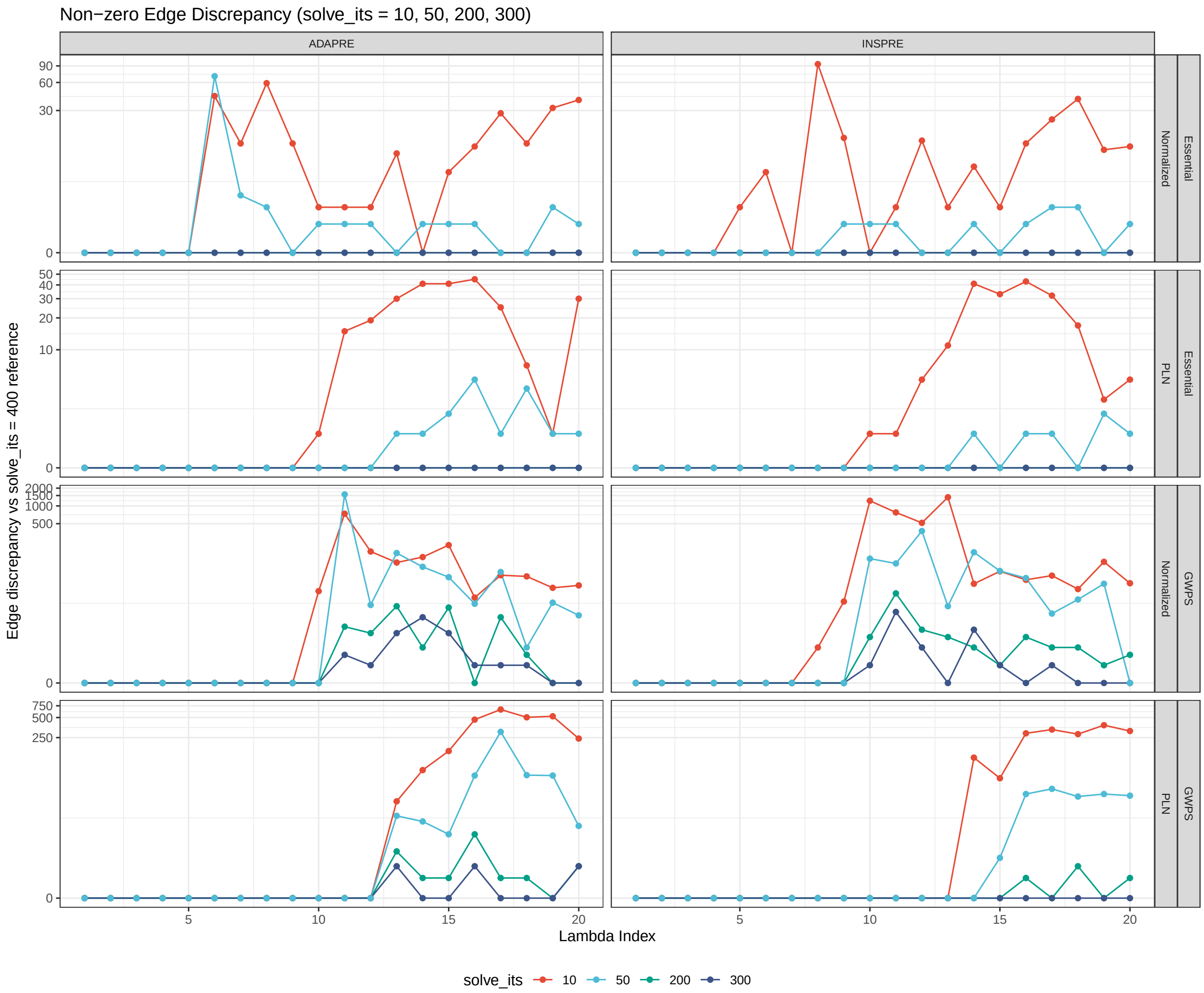


Figure S3: Stability of inferred sparsity with respect to `solve_its`. Discrepancy in the number of inferred nonzero edges at each $\lambda$ when using `solve_its`$\in \{10, 50, 200, 300\}$, relative to a high-accuracy reference fit with `solve_its`=400.

#### Supplementary Note S3: Directed precision against TF-binding references

We complement the enrichment analysis in the main text with a precision-oriented summary for directed TF→target edges. For each sparsity level, we compute the baseline-adjusted macro precision ΔPrec@50 (Variant B) against ChIP–seq TF-binding references (promoter, enhancer-linked, and all peaks).

Let $\widehat{B}$ denote an estimated directed-effect matrix at a given penalty level, and fix a threshold `thr` used to define predicted edges. For a transcription factor $t$, the predicted TF→gene edges are $\{(t, g) : |\widehat{B}_{tg}| > \texttt{thr}\}$, excluding the self-loop $t \to t$ when $t$ is in the evaluation gene set. We restrict evaluation to TFs with at least one reference-positive target ($n_{\text{ref}}(t) > 0$) and at least $K = 50$ predicted edges ($n_{\text{pred}}(t) \geq 50$). For each eligible TF $t$, we rank predicted targets by $|\widehat{B}_{tg}|$ and compute

$$\text{Prec@50}(t) = \frac{\#\{\text{reference-positive targets among the top 50}\}}{50}.$$

We report MacroPrec@50, the average of Prec@50($t$) over eligible TFs. To account for TF-specific reference density, we define a baseline binding rate for each eligible TF $t$ as $n_{\text{ref}}(t)/|G_t|$, where $G_t$ is the gene universe used for evaluation (excluding $t$ if $t \in G_t$). The final metric is

$$\Delta\text{Prec@50} = \text{MacroPrec@50} - \frac{1}{|\mathcal{T}|} \sum_{t \in \mathcal{T}} \frac{n_{\text{ref}}(t)}{|G_t|},$$

where $\mathcal{T}$ is the set of eligible TFs.

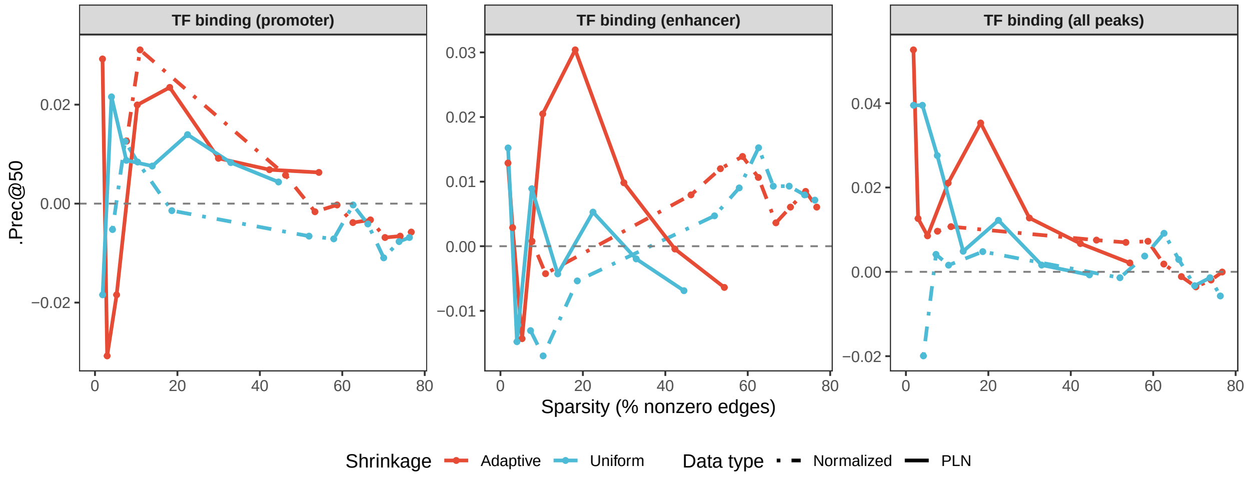


Figure S4: Baseline-adjusted directed precision against TF-binding references. ΔPrec@50 (Variant B) as a function of network sparsity for TF→target edges, evaluated against ChIP–seq TF-binding references constructed using promoter overlap, enhancer-linked overlap, and the union of all peaks. Curves compare uniform versus adaptive shrinkage and normalized versus PLN-based estimation.

#### Supplementary Note S4: Case-study comparison with *inspre*

We compared the K562 case-study GRNs inferred by ADAPRE and *inspre* across density-matched regularization paths on the common 300-gene universe.

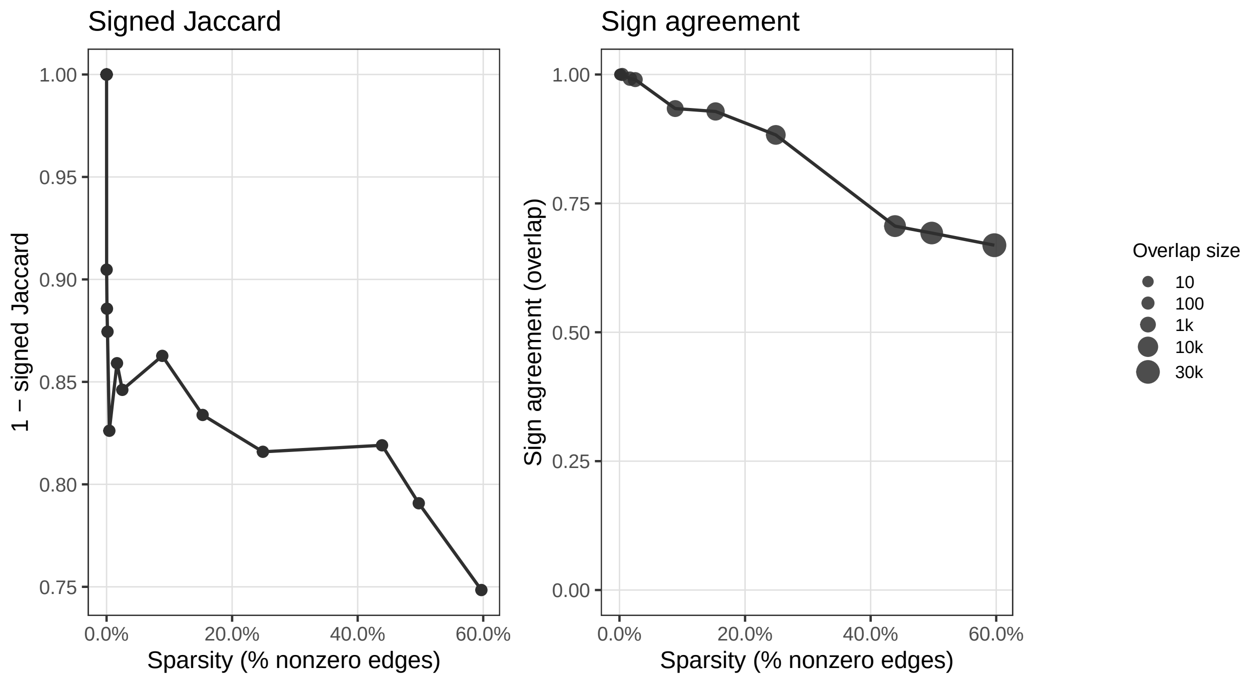


Figure S5: **ADAPRE vs. *inspre* case-study comparison on K562.** Across density-matched regularization paths on the common 300-gene universe (excluding self-loops), we report (left) signed Jaccard distance $1 - J^{\pm}$ between inferred directed networks (treating an edge with opposite sign as different) and (right) sign agreement among overlapping edges. Point size encodes the number of overlapping edges; at very sparse regimes overlap is near zero, inflating sign agreement toward 1 and motivating cautious interpretation in that region.
